# Video-Microscopy-Based Automated Trajectory Determination for High-Velocity, Densely Clustered, Indistinguishable Objects Moving in A Directed Force Field

**DOI:** 10.1101/2023.10.20.563243

**Authors:** Christopher Tyson, Santosh Gaire, Ian Pegg, Abhijit Sarkar

## Abstract

We present a method for tracking densely clustered, high-velocity, indistinguishable objects being spawned at a high rate and moving in a directed force field using only object centroids as inputs and no other image information. The algorithm places minimal restrictions on the velocities or accelerations of the objects being tracked and uses a methodology based on a scoring function and a back-tracking refinement process. This combination leads to successful tracking of hundreds of particles in challenging environments even when the displacement of the individual objects at successive times approaches the separation between neighboring objects in any one frame. We note that these cases can be particularly difficult to handle by existing methods. The performance of the algorithm is methodically examined by comparison to simulated trajectories which vary the temporal and spatial densities, velocities, and accelerations of the objects in motion, as well as the signal-to-noise ratio. Also, we demonstrate its capability by analyzing data from experiments with superparamagnetic microspheres moving in an inhomogeneous magnetic field in aqueous buffer at room temperature. Our method should be widely applicable since trajectory determination problems are ubiquitous in video microscopy applications in biology, materials science, physics, and engineering.

## Introduction

Automatic tracking of objects in images acquired through video microscopy is critical in many fields of engineering and science. The track estimation task in general involves two distinct steps (Thomann et al 2003). First, objects of interest - for example, particles, microspheres, cells, fluorophores - must be found and annotated in each image frame. Second, by comparing successive frames, trajectories for the annotated objects must be built up. The second task becomes easier if the objects are distinguishable, but the more challenging case is when the objects are indistinguishable. In this latter case, it is a priori not apparent how to map an object in one frame to itself in the next frame. If the objects are closely spaced, further complications arise since trajectory assignments for two neighboring objects may be switched by an algorithm without the altered trajectories deviating significantly from the true ones, making such errors especially hard to detect computationally.

A number of track determination algorithms have been developed and have performed with varying degrees of success, especially in the most challenging case of densely clustered, identical particles moving at high speed - see Meijering et al (2012) and Chenouard et al (2014) and references 32 – 57 therein. These algorithms may be divided into two categories: predictive tracking and measurement assignment tracking. In predictive tracking, image data are used to estimate an ensemble of kinematic models, and based on this ensemble, the algorithms determine probabilities for the current observations conditional on each possible trajectory in the ensemble. These probabilities can also be augmented with additional data – for example, Anderson et al use image intensity information (Anderson et al, 1992), while other methods have considered distributions of errors for observed parameters and kinetic model fits (Bar-Shalom and Tse, 1975). For densely clustered particles or significant noise, predictive tracking can offer an advantage. However, if the exact kinematic model is unknown or not accurately estimated – which can soften be the case – the effectiveness of these algorithms will suffer.

Measurement assignment tracking algorithms typically involve “scoring” each potential trajectory assignment and then using the scores to determine the best assignment. Velocities, trajectories, smoothness, shape, and size of objects of interest are often used to compute scores and must be estimated from the data. A number of techniques use a Kalman filter (Kalman, 1960) for this purpose (Cerveri et al, 2003). Constraints on the acceleration/deceleration, radius of turn, or inertia can be used to isolate only the objects of interest increasing computational efficiency (Barniv, 1985; Fortmann et al, 1983; Hashiro et al, 2002; Logothetis et al, 2002). However, one limitation is that these methods are primarily intended for tracking single objects under low noise conditions, although modifications exist to remove these constraints (Blanding et al, 2007; Chen and Tugnait, 2001; Hong et al, 1998). An alternative to Kalman-filter-based methods is the Multiple Hypothesis Tracking algorithm (Reid, 1979; Cox and Hingorani, 1996; Noyes and Atherton, 2004) in which all measurements prior to the current observations are compared against a predefined kinematic model to generate a set of parent hypotheses. Measurements in the current observation, as well as dummy measurements for noise, new objects, and false positives, are assigned a hypothesis with respect to the parent hypotheses, and Bayes theorem is used to calculate the probabilities of each current measurement based on the prior measurements.

These techniques, although powerful, tend not to perform well for densely clustered, indistinguishable particles moving unidirectionally at high speed and spawned at high rates, and, thus, new algorithms are called for. Here, we present an algorithm that fills the gap. Our method can be thought of as a hybrid. Its core is a measurement assignment tracking method – we do not use the data to fit an explicit kinematic model. However, it takes advantage of the underlying dynamics of objects moving in a unidirectional force field, much like a predictive tracking algorithm would, in order to provide an initial set of trajectories for the measurement-assignment-inspired step.

In the next section, we describe the algorithm in detail, as well as provide the procedures – simulation and experimental – used to validate it. This is followed by a detailed description of our results and a discussion of what implications they have for the accuracy and efficiency of our technique.

## Materials And Methods

### Algorithm

We wish to analyze scenes consisting of densely clustered, indistinguishable objects moving at high speed. These objects may be spawned at high rates, and their number frame-to-frame is not conserved as newly born objects add to the object population in successive frames and others exit the scene before traveling the entirety of the sensor’s field of view. Further, objects may have a variety of entrance points, may exit the field-of-view at any point for any reason, and they may have widely distributed velocities and other trajectory parameters.

The tracking algorithm involves two conceptually distinct but mutually supporting computations. The first involves a scoring function, which generates trajectory assignments for the particles, and the second involves the backtracking method, which takes as its input the scoring function output and further refines or modifies those trajectories. (The method presented here is independent of how particle centroid localization is performed.)

The scoring function, W_p,1;q,2_, calculates a score for each object p with coordinates (x_p,1_, y_p,1_) and at time 1 with respect to objects q at time 2 with coordinates (x_q,2_, y_q,2_), i.e., it assigns a real number to every possible pairing of objects in two consecutive frames. The score is not bounded, can take negative, positive, and zero values, and is defined as follows:

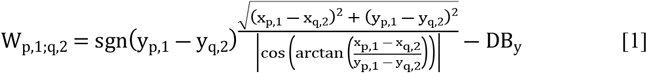

This can be simplified for computational purposes to read

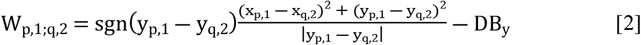

Referring to Equation 1, the first term in the scoring function is the signum function, which gives the score a positive value for movements in the direction of the force (which we remind the reader is taken to be in the negative-y direction) and a negative value for movements in the opposite direction. The numerator in Equation 1 is the Euclidean distance between the two observations while the denominator is the cosine of the angle formed between the direction of the force and the vector that connects the two objects. The final term, DB_y_, is calculated by finding the difference in the mean y-coordinates for all objects in two successive frames. However, since the location of a particle in the next frame has not been definitively assigned at this stage, it is calculated for different particle pairings and the minimum value is used. This term serves to penalize observation pairs which do not show substantial movement in the direction of the force and approximates the minimum distance an object is expected to move between successive observations based on the gross behavior of all objects.

The scoring function is constructed such that small, positive scores are preferred as these indicate movement in the direction of the force; large, negative scores, on the other hand, imply displacements in opposite or orthogonal directions.

The back-tracking approach provides a quality check for the score-based track assignments. It builds on the idea that if an object is observed at (x_1_, y_1_) at time 1 and subsequently at (x_3_, y_3_) at 3, then it is likely that for an observation at an intermediate time 2, the coordinates (x_2_, y_2_) should fall close to the straight line connecting (x_1_, y_1_) and (x_3_, y_3_). This is accomplished by calculating the following quantity:

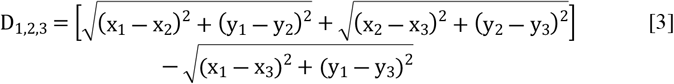

D_1,2,3_ is a measure associated with a frame triplet. It calculates the sum of distances from an object’s intermediate position (at time 2) to its positions at times 1 and 3 and from this subtracts the distance between its positions at times 1 and 3. Larger D-values indicate a significant deviation from the direct path between times 1 and 3, while smaller values represent closer alignment with the path; the limit of D = 0 represents the case of three collinear points.

Further discussion of the various aspects of the algorithm and its implementation, including a simplified illustrative example, can be found in the supplementary materials.

### Particle Trajectory Simulations

Performance of the algorithm was tested against simulated trajectories generated using the following inputs: (a) the total number of objects generated over the course of a simulation n_p_, (b) the size of the “image” in the x- and y-dimensions, (c) the spawn rate, the typical time between successive objects entering the field of view, (d) initial position (x_0_, y_0_), (e) initial velocity (v_x0_, v_y0_), (f) acceleration (a_y_) of each object, and noise components drawn independently for x- and y-object coordinates from a Normal distribution with mean zero and (g) variance σ, a measure of the signal-to-noise ratio. x_0_ is defined as the row coordinate of the pixel where a particle enters the simulated field of view; y_0,_ the column coordinate. The inputs can be single-valued or defined from an interval from which they may be drawn with uniform probability. The latter allows us to assess how well the algorithm can deal with objects with non-uniform behavior and also perform sensitivity analyses. The simulations continue until the final object exits the field-of-view of the “image.” Object coordinates are determined using the basic kinematical equations.

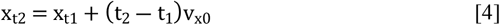

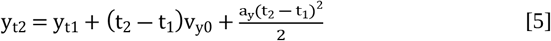

Here, (x_t1_, y_t1_) is the position of the particle at time t_1_ and (x_t2_, y_t2_) is its position at t_2_. Position is measured in pixels, velocity in pixels/frame, and acceleration in pixels/frame^2^. Given a specific pixel size and frame rate, these values can be converted to physical units. As the time slices are uniformly separated, if an object spawns at a time between two-time steps, its motion to the next time step is determined using the kinematical equations. Subsequently, it is simply a matter of stepping through time to calculate the spatial coordinates of the trajectory for each object. Noise is then added, and the final output is a list of observations (x, y, t) and trajectory ID numbers for each observation. We selected simulation parameters to cover a range of cases that might exist in experiments or data processing tasks. For further discussion on the rationale for the selection of simulation parameters, please see supplementary materials.

### Quantifying the Algorithm’s Performance

Two criteria were used to compare the results of the algorithm to experimental data and the ground truth trajectories known from simulations. First, the correct number of individual observation links is determined by comparing the simulation particle coordinates at each time step (ground truth) to the coordinates assigned to the same particles by the algorithm. From this, we determine if a particle has been correctly linked to itself at the next time step, and the total number of correctly identified links is recorded as a percentage of the known links.

The second approach utilizes the Variation of Information (VI) metric. This compares two partitions F and G of a set A, where each partition consists of disjoint subsets F = {f_1_, f_2_,…, f_e_} and G = {g_1_, g_2_,…, g_f_}, by computing the following quantity:

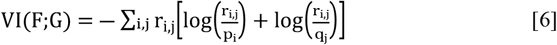

where 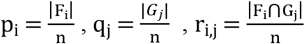, and n = Σ_i_ |F_i_| = Σ_j_ |*G*_*j*_| = |A|. In our case, the set A consists of all observations in the simulations. The partition F is the correct trajectory assignments from the simulations and the partition G is the algorithm output trajectories. The disjoint subsets within F and G are the individual object trajectories. As defined, VI is zero if the two partitions are identical. As the difference between the partitions grows, VI grows as well; thus, low values of VI are desirable.

### Experimental Validation

We performed experimental validation of our method by finding the magnetic force on magnetic microparticles moving in a buffer in a magnetic field by using to inferred trajectories of magnetic microparticles to compute the drag force and by a second independent method and comparing the two results. See supplementary materials for more details.

## Results

### Performance and Sensitivity Analyses on Simulation Data

Figures 1-5 display the algorithm’s performance varying each simulation parameter (excluding total particles) individually. Each figure has two columns with eight plots, illustrating the algorithm’s performance variation with changing parameters. The left column shows percent correct trajectory links (averaged over 50 simulations), while the right column presents VI scores, computed by grouping all 50 replications into a superset. Each column includes four plots, arranged by noise strength (0 to 3). The x-axis is spawn rate, and each plot represents specific spawn rates with grouped bars of different shades of gray, indicating the varied parameter. See the legend for an explanation of the gray shade values.

**Fig. 1.**
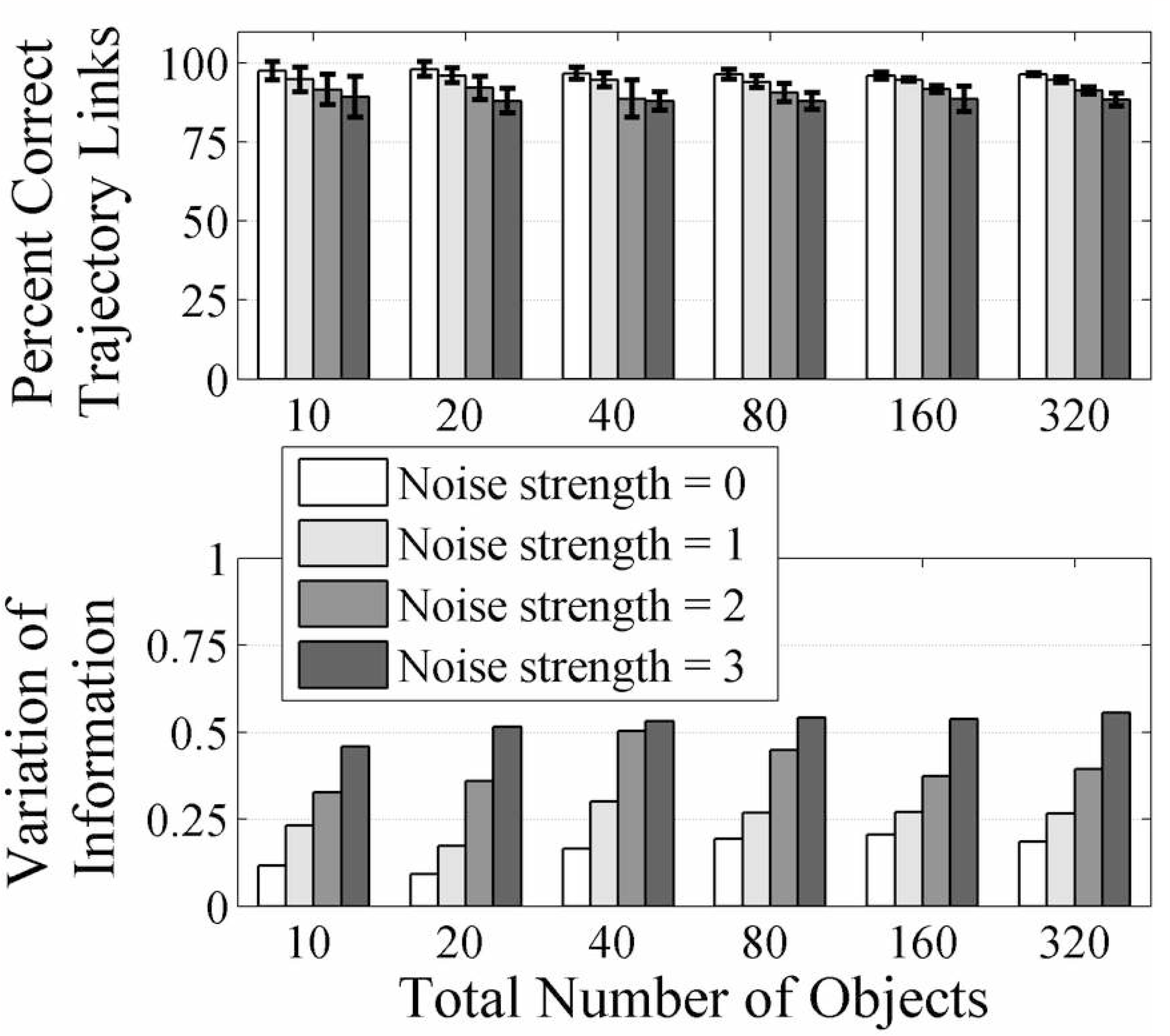
Results of simulations in which the number of total objects simulated was varied. All other simulation parameters remained in the baseline configuration with x _0_ = [1, 400], y_0_ = [1, 1], v_x0_ = [0, 0], v_y0_ = [0, 0], a_y_ = [9, 9] and spawn rate of 3/10, corresponding to 10 new objects every 3 frames. The simulations were run with added noise of strengths 0, 1, 2, and 3 as indicated by the legend.

We varied the total number of particles tracked [10, 20, 40, 80, 160, 320] in simulations with a spawn rate of 10/3 and other parameters at baseline values - see Fig.1. The algorithm showed excellent performance with close to 100% correct links for all particle numbers tested. Results were weakly sensitive to noise, dropping to about 90% at noise level 3 for all n_p_. Measured by VI, the performance degradation was expected but generally small (VI stayed below 1 for all noise and particle number values). However, for noise levels 1 and 2, VI behaved non-monotonically with particle numbers, increasing from 20 to 40, decreasing somewhat from 40 to 160, and rising again at 320.

In Fig. 2, we varied x_0_ by drawing from intervals of different lengths [200, 200], [150, 250], [100, 300], [50, 350], and [1, 400]. Optimal performance occurs when x_0_ is drawn from a wide distribution and the spawn rate is small (rate of 1 or lower). At rates between 2 and 8, wider sampling in x_0_ leads to better performance; for instance, for a rate of 4, the percentage of correct links varies from 70% to 95% as x_0_ is sampled from intervals of length 0 to 400. VI results align with the percent link findings. For high spawn rates and point sources, the VI is close to 3. However, without noise, performance significantly improves for rates of 8 or less, with VI less than 1, and almost 0 for very low spawn rates. The trend remains consistent with the introduction of noise: performance improves the most as x_0_ is drawn from longer intervals, approaching values with zero noise. Overall, we found weak sensitivity to noise across the parameter sets tested.

**Fig. 2.**
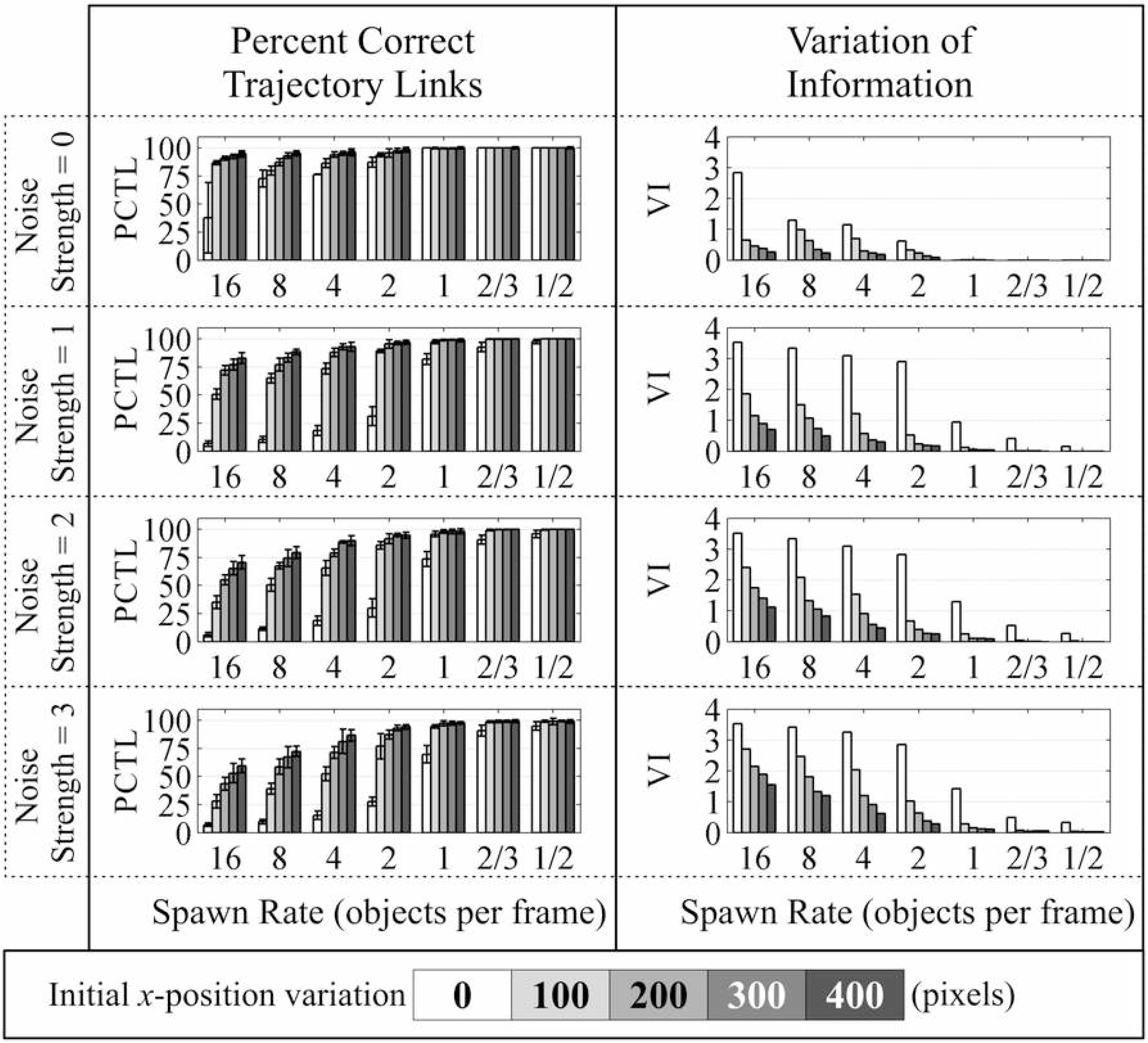
Results from simulations examining the initial x-position variation. While initial x-position variation was tested with values of 0, 100, 200, 300, and 400 (units of pixels), the remaining simulation parameters were held in the baseline configuration with n_p_ = 30, y_0_ = [1, 1], v_x0_ = [0, 0], v_y0_ = [0, 0], and a_y_ = [9, 9]. The two columns show the two-performance metrics: percent correct trajectory links in the left column, variation of information in the right column. Four rows correspond to the four different noise strengths: 0, 1, 2, and 3. The x-axis for all plots is the spawn rate in units of objects/frame. Each spawn rate grouping contains 5 bars of different shades of gray which represent the different initial x-position variation as shown in the legend at the bottom.

In Fig. 3, we varied initial y-position intervals [0, 50, 100, 150, 200]. Overall, the algorithm performed very well, with close to 100% correct identification and VIs less than 1 across a wide range of noise, spawn rates, and y_0_ interval lengths. However, performance declined as the interval length increased, modulated by the spawn rate, and showed low sensitivity to noise when spawn rate and interval length were constant. At high noise (strength 3) and spawn rates (8 and 16), performance dropped to below 50% for trajectory link identification, especially with higher y-position variation. Noise strength had limited impact on performance, except for specific cases like spawn rate 2, where VI slightly increased from 0.1 to 0.25 at interval length 0 for noise strength 3. At lower spawn rates, the differences in VI for various noise values were negligible.

**Fig. 3.**
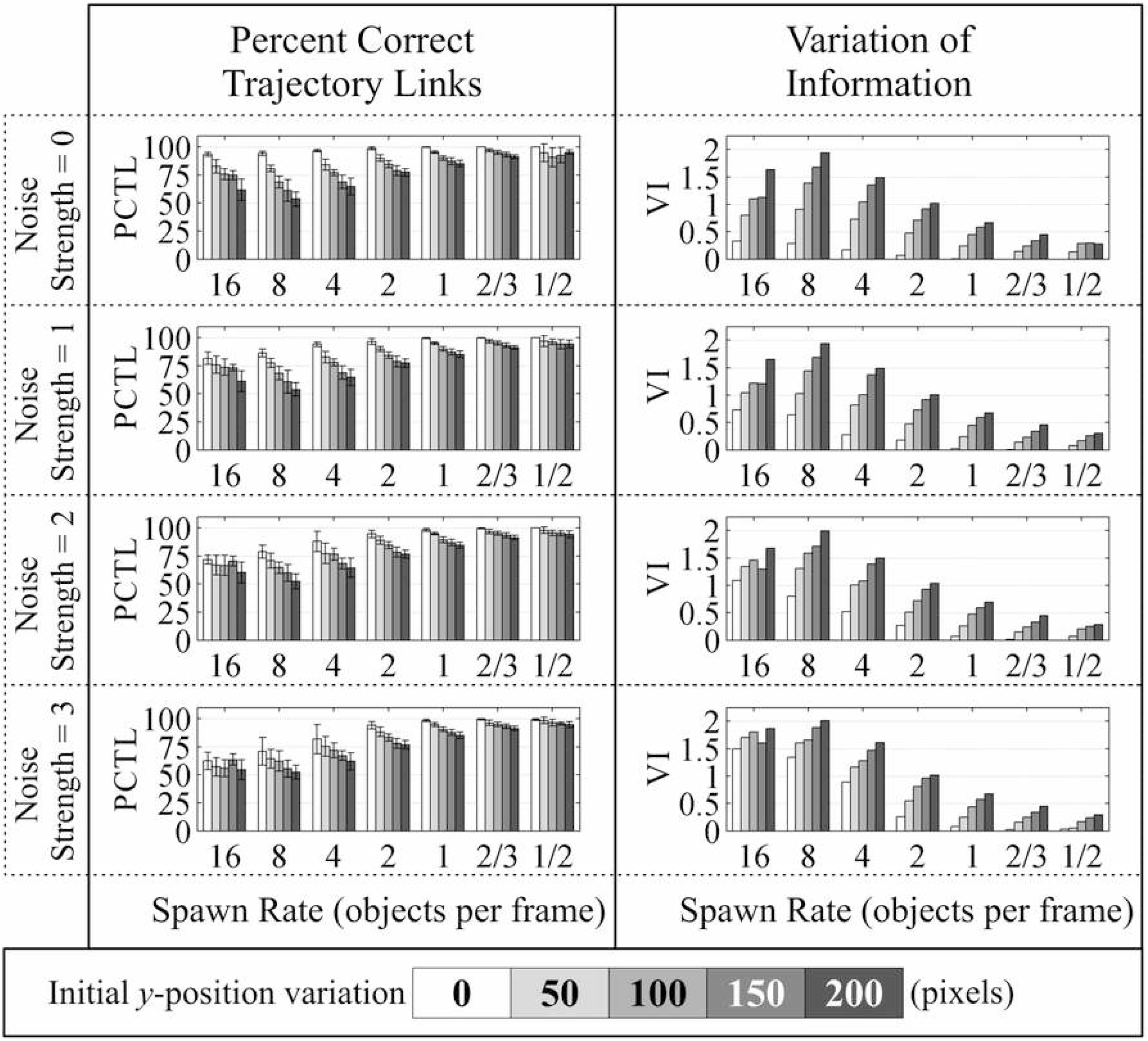
Results from simulations examining the initial y-position variation. While initial y-position variation was tested with values of 0, 50, 100, 150, and 200 (units of pixels), the remaining simulation parameters were held in the baseline configuration with n_p_ = 30, x_0_ = [1, 400], v_x0_ = [0, 0], v_y0_ = [0, 0], and a_y_ = [9, 9]. The two columns show the two-performance metrics: percent correct trajectory links in the left column, variation of information in the right column. Four rows correspond to the four different noise strengths: 0, 1, 2, and 3. The x-axis for all plots is the spawn rate in units of objects/frame. Each spawn rate grouping contains 5 bars of different shades of gray which represent the different initial y-position variation as shown in the legend at the bottom.

In Fig. 4, we drew the initial x-velocity (v_x0_) from varying intervals widths: [0, 0]; [-1, 1]; [-2, 2]; [-3, 3]; [-4, 4], corresponding to variations of 0, 2, 4, 6, and 8, respectively. An increase in variation generally led to decreased algorithm performance for fixed noise and spawn rates. For instance, at noise level zero and spawn rate 16, the percent correct trajectory links decreased from 95% to 75%, and VI increased from under 0.5 to close to 1.5 as the interval widened. However, with lower spawn rates (e.g., 2 or less), the overall performance improved significantly, nearing 100% correct identification and VI close to zero. These results remained robust even with increased noise amplitude. The worst performance occurred at spawn rate 1 and noise 3, but even then, over 95% of trajectory links were correctly identified, and VI remained below 0.25.

**Fig. 4.**
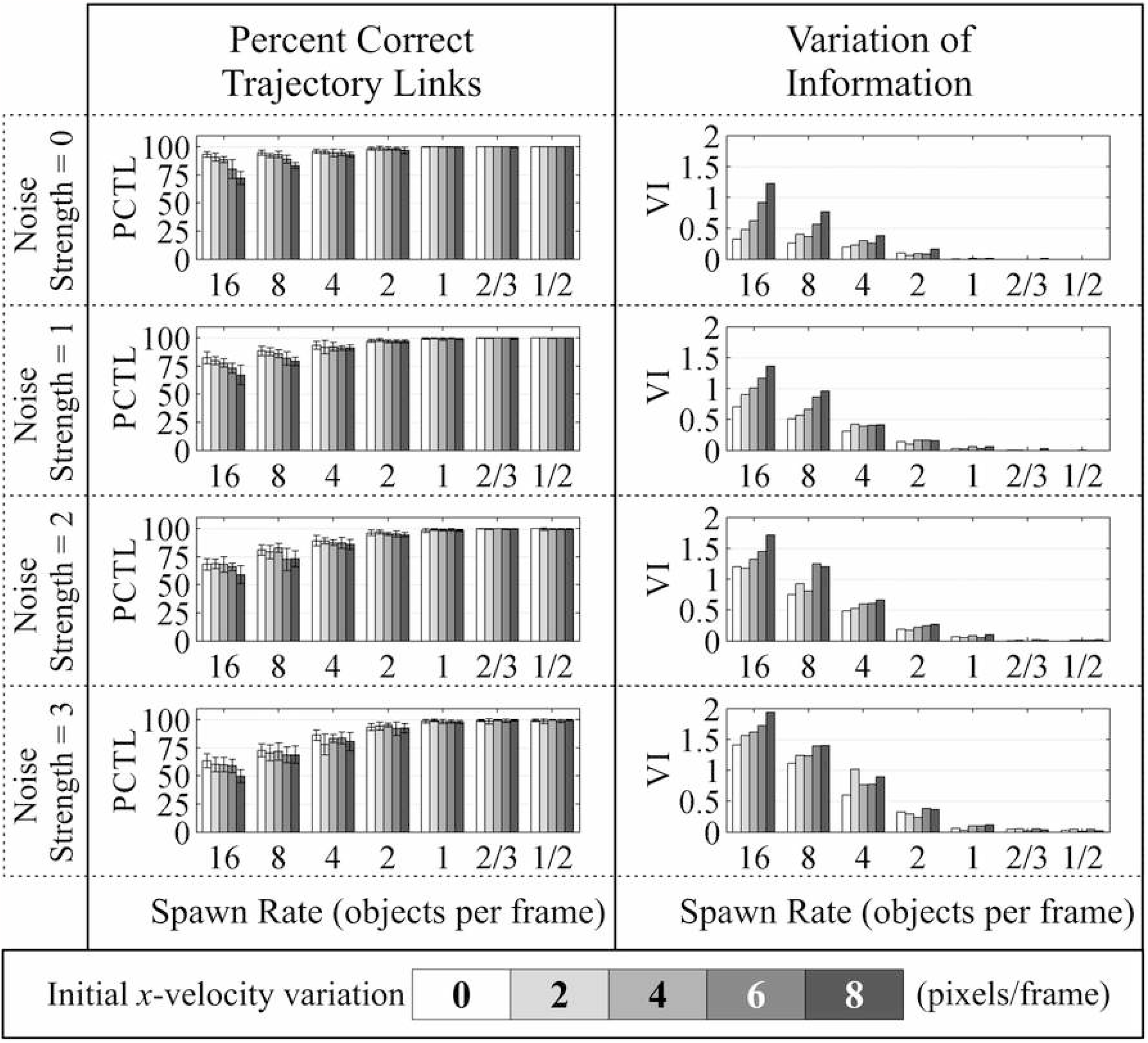
Results from simulations examining the initial x-velocity variation. While initial x-velocity variation was tested with values of 0, 2, 4, 6, and 8 (units of pixels/frame), the remaining simulation parameters were held in the baseline configuration with n_p_ = 30, x_0_ = [1, 400], y_0_ = [1, 1], v_y0_ = [0, 0], and a_y_ = [9, 9]. The two columns show the two-performance metrics: percent correct trajectory links in the left column, variation of information in the right column. Four rows correspond to the four different noise strengths: 0, 1, 2, and 3. The x-axis for all plots is the spawn rate in units of objects/frame. Each spawn rate grouping contains 5 bars of different shades of gray which represent the different initial x-velocity variation as shown in the legend at the bottom.

In Fig. 5, simulations were done with v_y0_ selected from intervals: [0, 0]; [0, 2]; [0, 4]; [0, 6]; [0, 8], representing variation lengths of 0, 2, 4, 6, and 8 respectively. At low noise strengths (0 and 1), a slight decrease in algorithm performance was observed as initial y-velocity variation increased. However, at higher noise strengths (2 and 3), the performance difference between y-velocity variations disappeared. The percent correct trajectory links drops from the mid-90% range to the mid-80% range for spawn rates of 16, 8, and 4. Vi measures did not show a discernible trend across spawn rates for noise strengths of 2 and 3. In such cases, the alterations caused by high noise strengths overwhelmed any impact of initial y-velocity variations.

**Fig. 5.**
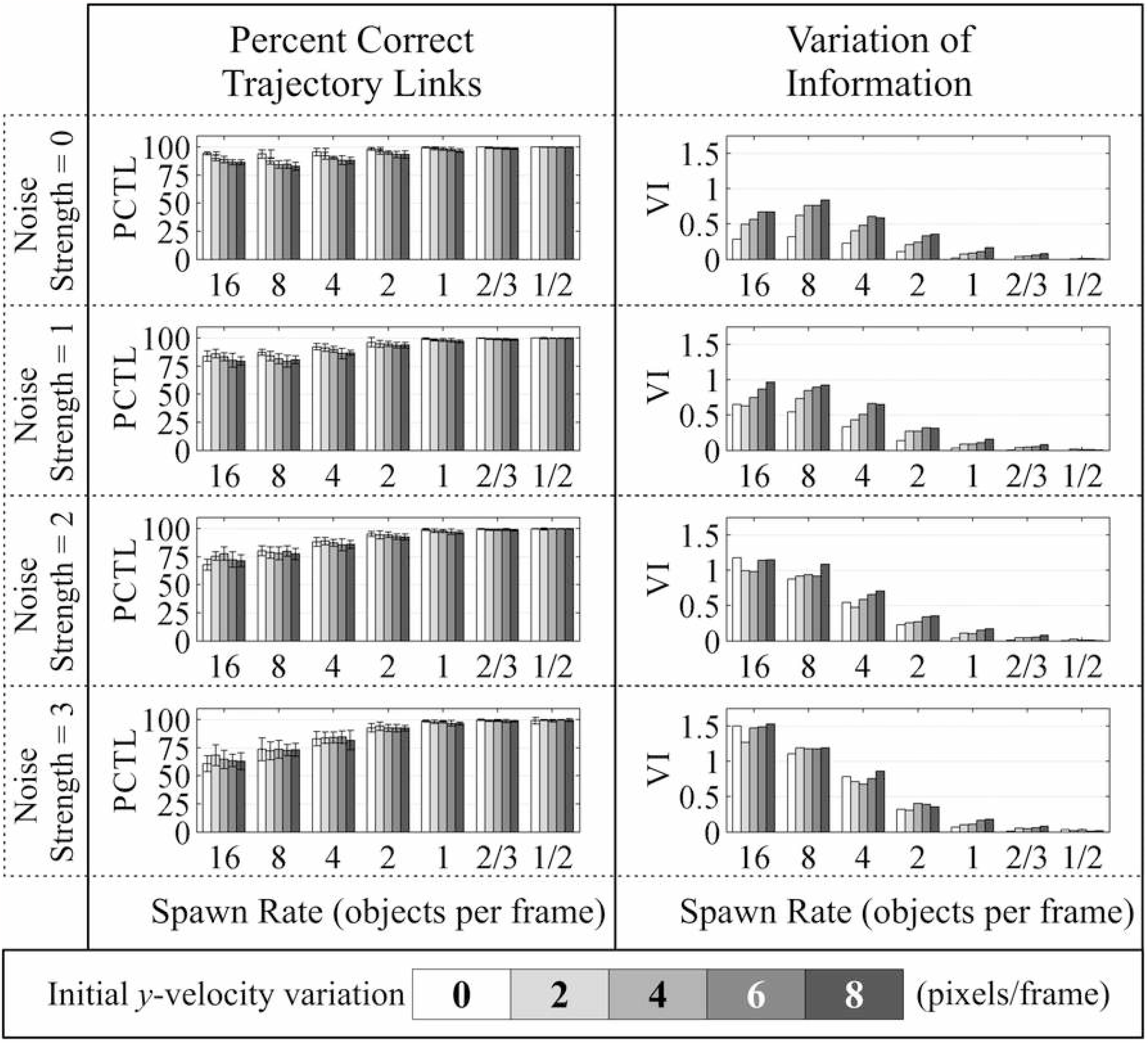
Results from simulations examining the initial y-velocity. While initial y-velocity variation was tested with values of 0, 2, 4, 6, and 8 (units of pixels/frame), the remaining simulation parameters were held in the baseline configuration with n_p_ = 30, x_0_ = [1, 400], y_0_ = [1, 1], v_x0_ = [0, 0], and a_y_ = [9, 9]. The two columns show the two-performance metrics: percent correct trajectory links in the left column, variation of information in the right column. Four rows correspond to the four different noise strengths: 0, 1, 2, and 3. The x-axis for all plots is the spawn rate in units of objects/frame. Each spawn rate grouping contains 5 bars of different shades of gray which represent the different initial y-velocity variation as shown in the legend at the bottom.

We performed additional simulations with altered initial y-acceleration: a_y_ = [9, 9]; [7, 11]; [5, 13]; [3, 15]; [1, 17], representing variation in values of 0, 4, 8, 12, and 16 respectively – see Fig. S5 in supplementary materials. For zero noise, a y-acceleration variation of 0 led to notably better algorithm performance in both percent correct trajectory links and VI. However, for y-acceleration variations greater than 0, there was no clear trend in performance. The differences in VI values were less noticeable at higher noise strengths, and in some cases, VI for y-acceleration variation of 0 was greater than the maximum variation of 16. When the noise strength is 3 and the spawn rate is 16, we see that VI for y-acceleration variation of 0 is greater than when the y-acceleration variation is at the maximum, 16. Overall, there was a slight improvement in performance with low y-acceleration variation, but the difference was often minimal and within the standard deviation of other values.

### Experimental Tests

We also carried out experimental tests to assess the accuracy of our method by releasing neutrally buoyant, micron-sized magnetic beads suspended in buffer in the magnetic field of a small bar magnet. In the low Reynolds number condition of the experiment, the drag force, which is proportional to the velocity, is equal to magnetic force. By computing the magnetic force in two independent ways, we were able to compare the particle-trajectory-based method to a second, orthogonal method. we found good agreement between the velocity-based method, which used the particle tracking algorithm to determine velocity, and the reference method.

Please see supplementary materials for simulation studies designed to recapitulate experimental conditions, details on the experimental tests, as well as studies on algorithm run time.

## Discussion

We have presented an algorithm for determining trajectories of closely spaced, indistinguishable objects that are spawned at high rates and are moving rapidly in a directional force field. The algorithm combines a scoring function that considers expected motion due to the force field with a back-tracking method inspired by measurement-assignment techniques. In order to test its performance, we carried out simulation-based validation and sensitivity studies in which we systematically varied the spawn rate, initial conditions for object position and velocity (x_0_, y_0_, v_x0_, v_y0_), object acceleration (a_y_), and noise strength or the signal-to-noise ratio (σ). We also looked at how the performance scaled with n_p_, the total number of particles tracked. These were complemented by additional simulations designed to recapitulate microsphere tracking experiments (described in supplementary materials) and experimental validation was performed as well. Results were quantified in terms of (a) the percentage of correct links identified and (b) the Variation of Information (VI) score.

Starting with n_p_, we found a weak dependence on particle number. This is not surprising since we expect entry of new objects to be compensated to some extent by the exit of others, roughly leaving the same number of particles to be tracked in each frame independent of the total number of particles.

The performance was generally robust as a function of the initial kinematical parameters of the particles (x_0_, y_0_, v_x0_, v_y0,_ a_y_) with successful link detection of 90% or higher over a wide range of values for these quantities, even for objects with large variations in velocities or accelerations (i.e., when, in simulations, we drew these values from intervals of varying lengths.) Also, we found that the spawn rate and, to a lesser extent, the noise strength played a bigger role in limiting the algorithm.

As x_0_ is drawn from wider intervals, the resulting trajectories tended to be laterally spaced apart, which helped with the identification task as misidentifications would lead to an object in the next frame undergoing a large lateral displacement, an assignment that was heavily penalized by the scoring function. Conversely, a point source leads to tightly clustered paths (the location of the point source does not matter – data not shown) and this case is difficult to parse. This is because the lateral separation between links may only be a few pixels making scoring-based discrimination less effective. This also explains why the performance measured in VI for the point source case is very sensitive to noise. Even for zero noise, trajectories are packed together tightly, leading to a challenging discrimination task. When noise is added, the chances of erroneous link assignments go up even more making it that much harder to assign true trajectories. For y_0_, at high spawn rates, increasing the length of the interval increases the probability that a new object A will spawn at time t_j_ very close to the position of object B at time t_i_. If A is closer to B’s location at t_j-1_ than B at t_j_ is to its previous value, the algorithm will determine that A should be assigned to the trajectory of object B, while assigning B at t_j_ to a new trajectory, leading to link (and trajectory) misidentifications.

When we looked at the effect of increasing the variation in v_x0_ we expected improved performance since greater non-uniformity in v_x0_ can increase horizontal separation between objects. However, because we allow objects to have both positive and negative x-velocity, the chance of multiple path intersections increases. The algorithm is capable of dealing with path intersections when the trajectories are spaced in time adequately (lower spawn rates) but as the objects are grouped closer and closer together (at high spawn rates) it can become quite challenging to discern between trajectories that intersect frequently within a small space. These trends were recapitulated for v_y0_ where an increase in v_y0_ led to a slight decrease in performance. As the scoring function penalizes motion against the direction of the force, variations in the movement of objects in the direction of the force will see a smaller change in algorithm performance. As for a_y_, the results were robust to non-uniformities in the acceleration of the objects.

When looking at how performance depends on noise and spawn rate, in general we found a more prominent role for the spawn rate, although the signal-to-noise ratio affected results as expected. Indeed, increasing noise strength correlates with a decrease in performance. This is especially true for trajectories that are closely spaced since densely clustered objects can essentially swap positions and still have trajectories close to the true ones. This is evident in Fig. 2, which plots results from altering the initial x-position variation. In these simulations, the most densely clustered objects are represented by an initial x-position variation of 0 and a spawn rate of 16, with 16 objects appearing from the same exact point in every frame. As we look at the change in performance metrics from noise strength of 0 to noise strength of 1, we see a significant decrease in performance with VI values increasing 2-fold for noise strength of 1 compared to noise strength of 0. As the objects increase in separation (the initial x-position variation is larger), the effect of noise on the algorithm performance is less profound. Furthermore, when the objects are separated both spatially and temporally, the noise strength has even less influence on the algorithm performance. Again, referring to Fig. 2, initial x-position variations of 300 or 400 and spawn rates of 1 or less lead to barely noticeable changes in VI values.

One of the fundamental limits of tracking algorithms is how closely the objects are located to one another. This is a function not only of the objects’ motion in space, but also of the rate at which they enter or exit the sensor’s field of view, i.e., the rate at which they spawn. The role of the objects’ spatial density was evident in our simulations when we studied how sensitive the algorithm was to variations in the initial positions, velocities, and accelerations. These three parameters were found to have the most significant effect on object density with greater variation in their values leading to greater spatial separation. (The caveat is that greater variation in x-velocity and y-acceleration can also increase the frequency of trajectory intersections.)

We determined mean nearest neighbor distances for objects in these trials, i.e. the mean distance between each object at time t and its nearest neighbor at that same time. Deciding between the nearest neighbor and the true object is the core challenge that the algorithm faces. These nearest neighbor distances vary depending on the simulation parameters used, including both spatial parameters and spawn rates. At a spawn rate of 16, the mean nearest neighbor distance is between 3 and 19 pixels, regardless of the spatial parameters. Reducing the spawn rate to 4, the mean nearest neighbor distance grows to between 4 and 50 pixels. Then at a spawn rate of 1, the mean nearest neighbor distance spans the range of 40 to 140 for all spatial parameters. It is interesting to note that we see instances in the simulations where high spawn rate (>=8) and dense object clustering can often present nearest neighbor distances of less than a single pixel, and sometimes even with multiple object centroids within a single pixel. Parsing trajectory assignments for objects spaced so closely is incredibly challenging, particularly so without utilizing any special image processing methodology.

The spawn rate also played a significant role in the ability of the algorithm to discriminate trajectories. Indeed, in any of the Figs. 2-5, an examination of the same color bars of any sub-plot shows that the performance of the algorithm improves consistently as the object spawn rate is decreased. For object spawn rates of 1 or lower, the worst performance of the algorithm across all variables occurs for at an initial x-position variation of 0 and noise strength of 3, in which 70% of trajectory links were correctly identified and a VI of 1.4 is achieved. This is perhaps one of the most difficult cases to parse accurately, and yet the algorithm is capable of detecting 70% of trajectory links correctly while maintaining a VI of 1.4. In fact, the only other case in which VI is greater than 1 for a spawn rate of one is when the initial x-position variation is 0 and the noise strength is 2. For all other parameter value combinations, VI is below 1. Even for a spawn rate of 2, the algorithm only has significant issues when the initial x-position variation is 0; for all other parameter combinations, VI at spawn rates of 2 does not significantly exceed 1. In an experimental setting, this means that if the frequency of observation (frame rate) is high enough to reduce the spawn rate to some value around 2, the algorithm should have excellent performance regardless of object dynamics.

The backtracking method was evaluated by comparing correct trajectory links of the scoring function to those of the complete algorithm. For the set of simulation parameters discussed previously, the number of correct trajectory links using the entire algorithm were compared to the number of correct trajectory links using the scoring function only, and the difference between the two values was calculated – see Fig. 6. In this figure, histogram bins less than zero represent trials in which the back-tracking method resulted in a decrease in the number of correct trajectory links, while positive bins count simulations where an improvement after back-tracking was achieved. We see that while there are situations in which the back-tracking method can reduce the effectiveness of the scoring function-based trajectory assignment, most often the back-tracking method improves upon the initial trajectory assignments. We also note that for the bulk of the simulations, the back-tracking method makes in the range of 0-25 trajectory assignment corrections after examining hundreds of inputs. This suggests that the scoring function is likely capable of performing rather well on its own, but an improvement on the scoring function is seen in the majority of cases after passing through the backtracking method.

**Fig. 6.**
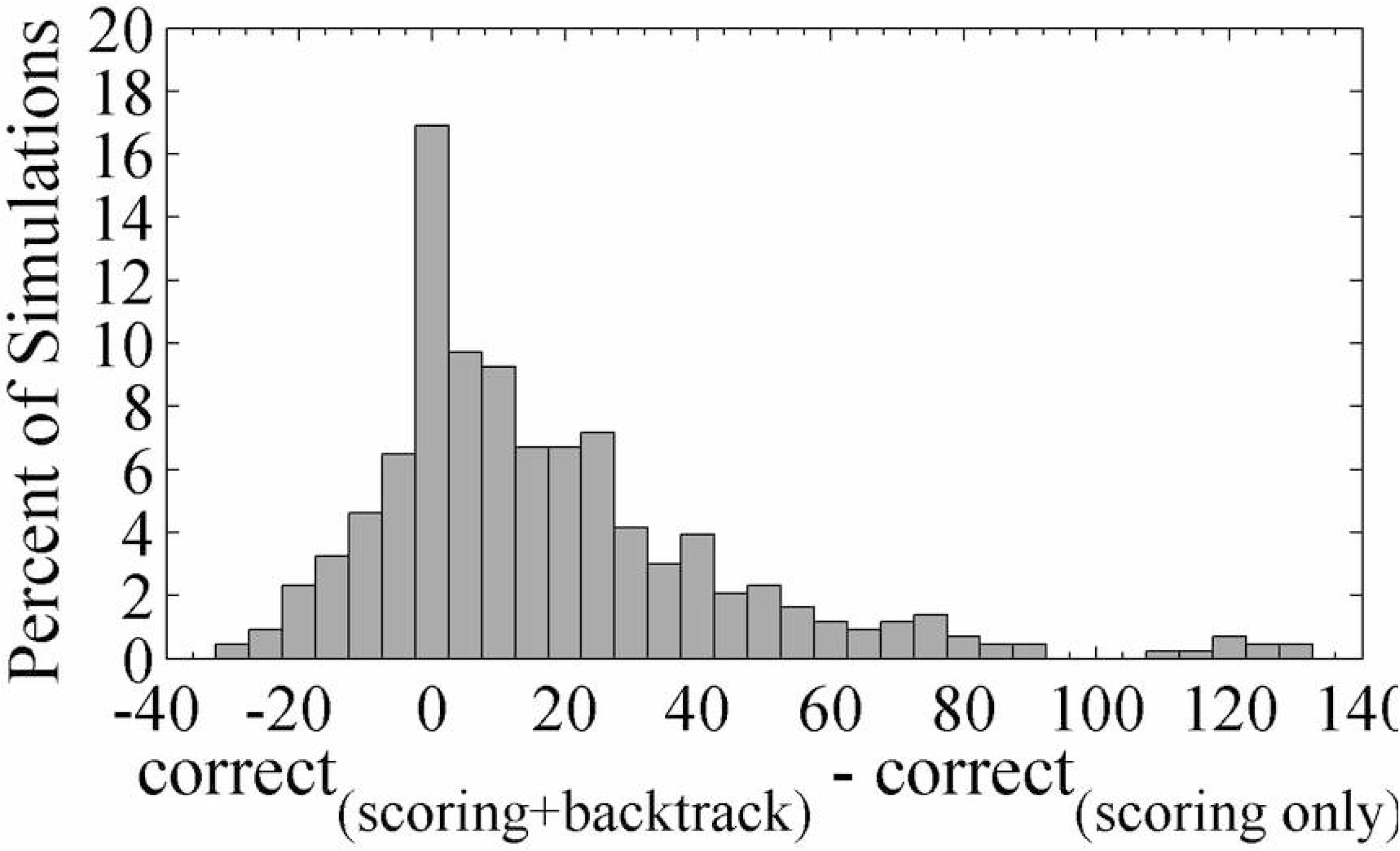
Examining the effect of the back-tracking method on the scoring function output. Here we show how the back-tracking method may alter the scoring function output from the set of simulation trials shown previously. The x-axis values correspond to results that were calculated by subtracting the number of correct assignments from the scoring function only from the number of correct trajectory assignments using both the scoring function and back-tracking method. Thus, a positive number indicates trials which saw improvement with the back-tracking method, while negative numbers correspond to trials in which the back-tracking method caused additional incorrect assignments. These results were collected into bins of width 5, and so each bar shows the percentage of the total number of trials in that bin. The bin values are half-closed intervals, e.g. [0,5) for the bin labeled 0, or [5,10) for the bin labeled 5.

Finally, we carried out experimental tests to assess the accuracy of the algorithm by comparing the forces imputed by the algorithm with those determined independently using an orthogonal comparator method. As discussed in the supplementary materials, we found good agreement between the algorithm-based and reference methods.

## Acknowledgements

We gratefully acknowledge the assistance of the following individuals: German Uritsky, Kara Fagerstrom, Josh Maher, Dunja Skoko, C.L. Chou, Eric Fischer, Robert Mohr, and Roberto Fabian. A. Sarkar would like to acknowledge funding from Vitreous State Laboratory and The Catholic University of America.

## Author Contributions

CT and SG programmed the algorithm and simulations; CT and SG carried out the experiments and related data analysis; CT and AS designed the algorithm and simulations; ILP, and AS designed and supervised the experiments and assisted with the data analysis; CT, SG, ILP, and AS wrote the manuscript.

## References

Anderson CM, Georgiou GN, Morrison IEG, Stevenson GVW, Cherry RJ. Tracking of cell surface receptors by fluorescence digital imaging microscopy using a charge-coupled device camera. 1992. Low-density lipoprotein and influenza virus receptor mobility at 4 degrees C. J Cell Sci 101: 415–425.

Barniv Y. Dynamic programming solution for detecting dim moving targets. 1985. IEEE Trans Aerosp Electron Syst 21: 144–156. doi: 10.1109/TAES.1985.310548.

Bar-Shalom Y, Tse E. Tracking in a cluttered environment with probabilistic data association. 1975. Automatica 11: 451–460. doi: 10.1016/0005-1098(75)90021-7.

Blanding WR, Willett PK, Bar-Shalom Y. 2007. Offline and real-time methods for ML-PDA track validation. IEEE Trans Signal Process 55: 1994–2006. doi: 10.1109/TSP.2007.893212.

Cerveri P, Pedotti A, Ferrigno G. 2003. Robust recovery of human motion from video using Kalman filters and virtual humans. Hum Mov Sci 22: 377–404. doi:10.1016/S0167-9457(03)00004-6.

Chen B, Tugnait JK. 2001. Tracking of multiple maneuvering targets in clutter using IMM/JPDA filtering and fixed-lag smoothing. Automatica 37: 239–249. doi:10.1016/S0005-1098(00)00158-8.

Chenouard N, Smal I, Chaumont FD, Maska M, Sbalzarini I, Gong Y, et al. 2014. Objective comparison of particle tracking methods. Nature Methods 11: 281–289. doi:10.1038/nmeth.2808.

Cox II, Hingorani SL. 1996. An efficient implementation of Reid’s multiple hypothesis tracking algorithm and its evaluation for the purpose of visual tracking. IEEE Trans Pattern Anal Mach Intell 18: 138–150. doi: 10.1109/34.481539.

Fortmann TE, Bar-Shalom Y, Scheffe M. 1983. Sonar tracking of multiple targets using joint probabilistic data association. IEEE J Oceanic Eng 8: 173–184. doi: 10.1109/JOE.1983.1145560.

Hashiro M, Kawase T, Sasase I. Maneuver target tracking with an acceleration estimator using target past positions. 2002. Electro Commu Jpn Part I 85: 29–37. doi: 10.1002/ecja.10026.

Hong L, Cui N, Cong S, Wicker D. 1998. An interacting multipattern data association (IMPDA) tracking algorithm. Signal Processing 71: 55–77. doi:10.1016/S0165-1684(98)00134-0.

Kalman RE. 1960. A new approach to linear filtering and prediction problems. J Basic Eng 82: 35–45.

Logothetis A, Krishnamurthy V, Holst J. 2002. A Bayesian EM algorithm for optimal tracking of a maneuvering target in clutter. Signal Processing 82: 473–490. doi:10.1016/S0165-1684(01)00198-0

Meijering E, Dzyubachyk O, Smal I. 2012. Methods for cell and particle tracking. Methods Enzymol 504: 183–200.

Noyes SP, Atherton DP. 2004. Control of false track rate using multiple hypothesis confirmation, target tracking. Algorithms and Applications, IEEE 115–120. doi: 10.1049/ic:20040062.

Reid DB.1979. An Algorithm for Tracking Multiple Targets. IEEE Trans Automat Contr 24: 843–854. doi: 10.1109/TAC.1979.1102177.

Thomann D, Dorn J, Sorger PK, Danuser G. 2003. Automatic fluorescent tag localization II: Improvement in super-resolution by relative tracking. J Microsc 211: 230–248. doi: 10.1046/j.1365-2818.2003.01223.x.

